# Inhibition and down regulation of the serotonin transporter contribute to the progression of degenerative mitral regurgitation

**DOI:** 10.1101/2020.03.10.985382

**Authors:** Robert J Levy, Emmett Fitzpatrick, Estibaliz Castillero, Halley J Shukla, Vaishali V Inamdar, Arbi E Aghali, Juan B Grau, Nancy Rioux, Elisa Salvati, Samuel Keeney, Itzhak Nissim, Robert C Gorman, Lubica Rauova, Stanley J Stachelek, Chase Brown, Abba M Krieger, Giovanni Ferrari

**Author notes:** Address for Correspondence: Giovanni Ferrari PhD - Associate Professor of Surgery and Biomedical Engineering, Columbia University: 630W 168^th^ Street 17.401c New York, NY 10032, E.

## Abstract

**Aims:** Heart valve disease attributed to serotonin (5HT) has been observed with 5HT-secreting carcinoid tumors and in association with medications, such as the diet drug, Dexfenfluoramine, a serotonin transporter (SLC6A4) inhibitor and 5HT receptor (HTR) 2B agonist. HTR2B signaling upregulates TGFβ-1 resulting in increased production of extracellular matrix proteins. SLC6A4 internalizes 5HT, limiting HTR signaling. Selective 5HT reuptake inhibitors (SSRI), widely used antidepressants, target SLC6A4, thus enhancing HTR signaling. However, 5HT and SLC6A4 mechanisms have not been previously associated with degenerative mitral regurgitation (MR). The present studies investigated the hypothesis that both dysregulation of SLC6A4 and inhibition of SLC6A4 contribute to the pathophysiology of MR.

**Methods and Results:** Here we report SLC6A4 related studies of 225 patients with MR requiring surgery. A multivariate analysis showed that SSRI use in MR patients was associated with surgery at a younger age, indicating more rapidly progressive MR (p=0.0183); this was confirmed in a national dataset (p<0.001). Aspirin use by MR patients was associated with surgery at an older age (p=0.0055). Quantitative reverse transcriptase PCR of MR leaflet RNA from 44 patients, and 20 normal mitral leaflets from heart transplant recipients, demonstrated down regulation in MR of both SLC6A4 and vesicular monoamine transporter-2 (SLC18A2), that packages 5HT (p<0.001). Human mitral valve interstitial cells cultivated with Fluoxetine, a SSRI, demonstrated down regulation of SLC6A4 and upregulation of HTR2B, compared to untreated, in cells from both normal and MR leaflets. Platelet 5HT studies in healthy subjects without heart disease used ADP-induced activation to model MR-associated activation. Fluoxetine significantly increased platelet activation and plasma 5HT levels, while Aspirin inhibited ADP platelet activation.

**Conclusions:** Down regulation and inhibition of SLC6A4 influences MR through enhanced HTR signaling. SSRI may further influence MR through inhibition and down regulation of SLC6A4, upregulation of HTR2B, and increased platelet release of 5HT.

**Translational Perspective:** Degenerative mitral valve regurgitation (MR) affects millions, and there is no medical therapy for this disease. MR becomes progressively worse, and for severe MR, the only option is cardiac surgery. Serotonin (5HT) is best known as a neurotransmitter. However, 5HT secreting carcinoid tumors cause a cardiac valve disorder in many cases, and 5HT related medications, such as the diet drug Fenfluoramine, have been associated with the development of cardiac valve disease. The present paper presents evidence that diminished serotonin transporter (SLC6A4) expression and inhibition, lead to increased 5HT receptor signaling, contributing to the progression of MR.

## Introduction

Serotonin (5HT) related mechanisms have been associated with clinically significant cardiac valvulopathies occurring with either 5HT secreting carcinoid tumors ^1^, or the administration of serotonergic agents, such as the diet drug, Dexfenfluramine^2-4^, a serotonin transporter (SLC6A4) inhibitor ^5^. Dexfenfluoramine also has high affinity for the 5HT receptor 2B (HTR2B) ^6, 7^, and is hypothesized to be an agonist for this receptor ^7^. Dopaminergic agents, such as the anti-Parkinson’s agent, Pergolide ^8^, have also been associated with a valvulopathy attributed to HTR2B agonist activity ^6^. The mechanisms responsible for 5HT-valvulopathies are incompletely understood. Both HTR2A and HTR2B signaling have been demonstrated to upregulate TGFβ-1, that in turn increases extracellular matrix protein expression in heart valve interstitial cells ^9-12^ and fibrotic diseases ^13^. SLC6A4 is a transmembrane protein that processes 5HT intracellularly ^14-16^ thereby limiting HTR activity. SLC6A4 is the molecular target of the class of antidepressants known as selective serotonin reuptake inhibitors (SSRI) ^17^. SSRI use by cardiac valve disease patients has been previously studied, with publications both implicating SSRI ^18^ and others finding no relationship of SSRI use to heart valve disease ^19, 20^. SLC6A4 is also present in mitral valve interstitial cells (MVIC) ^21^. A mouse model with SLC6A4 deleted demonstrated increased thickening of the mitral and aortic valve leaflets, as well as myocardial fibrosis, compared to non-deleted controls ^22^, thus suggesting that diminished SLC6A4 expression plays a role in the pathophysiology of cardiac valve disease.

Degenerative mitral regurgitation (MR), due to myxomatous mitral valve disease ^23^, when severe can only be treated surgically ^24, 25^. There is little known about 5HT-mechanisms and MR. ^21, 26^. Previous investigations of 5HT-related gene expression changes in human MR leaflets were reported by our group in studies of leaflets from four MR cases compared to four normal mitral leaflet samples ^26^. This study reported significant down regulation of SLC6A4 in MR leaflets compared to normal mitral leaflets. It is also noteworthy that MR is associated with a chronic state of platelet activation ^27, 28^, that correlates with the severity of valvular regurgitation ^27^. Since platelets release 5HT when activated ^29^, MR patients over time would hypothetically have more 5HT exposure than subjects without MR. Aortic stenosis has also been shown to result in increased platelet activation, and elevated plasma 5HT levels as well ^30^. Thus, the present studies investigated the hypothesis that 5HT and related SLC6A4 mechanisms contribute to the pathophysiology of MR.

## Materials and Methods

### Patient enrollment (Table 1)

Patients with MR, who were referred for first time surgery at the participating hospitals from 2009-2017 were enrolled in this study. Informed consent per IRB approval was obtained at either The Hospital of the University of Pennsylvania (IRB Protocol #809349) or The Valley Hospital (IRB Protocol #11.0009), upon admission prior to surgery. All patient information was de-identified. Normal mitral valve leaflets, together with de-identified clinical data, were obtained from the heart transplant service of the Hospital of the University of Pennsylvania with IRB approval (IRB Protocol #802781). All human subjects’ research in this paper, including the use of human tissues, conformed to the principles outlined in the Declaration of Helsinki. Exclusion criteria for this study included Marfan’s syndrome, congenital mitral valve abnormalities, endocarditis, rheumatic heart disease, ischemic mitral regurgitation, a history of cancer, autoimmune diseases, previous mitral surgery, and any history of cardiac trauma. To the best of our knowledge no patients with the X-linked Filamin-A mutation associated with mitral regurgitation ^31^ were included. Normal leaflet retrievals also excluded valve leaflets from patients with mitral valve disease. SSRI use prior to surgery was ascertained by searching for the following drugs: Citalopram, Fluoxetine, Fluvoxamine, Paroxetine, Sertraline, and Vilazodone.

**Table 1:** Characteristics of the degenerative mitral regurgitation (MR) surgical patient population and transplant recipients providing normal mitral valve leaflets (means ± standard deviations)

| Parameter | Entire population | MR subjects (from the 225 entire population) with qRT-PCR leaflet analyses | Normal leaflet subjects (transplant recipients) with qRT-PCR analyses |
| --- | --- | --- | --- |
| **Total** | 225 | 44 | 20 |
| Men | 160 (71.1) | 31 (70.5) | 13 (65.0) |
| Women | 65 (28.9) | 13 (29.5) | 7 (35.0) |
| **Age at surgery, years (range)** | 24 - 89 | 31 - 89 | 21 - 69 |
| Mean age at surgery, years (± SD) | 63.1 ± 12.7 | 59.6 ± 16.5 | 51.7 ± 13.3 |
| Men, years | 62.9 ± 12.2 | 58.1 ± 15.9 | 51.5 ± 12.5 |
| Women, years | 63.8 ± 14.1 | 63.2 ± 17.9 | 52.0 ± 15.8 |
| **Caucasian** | 203 (90.2) | 41(93.2) | 14 (70.0) |
| **Mitral Valve (MV) Surgical Intervention** |  |  |  |
| MV Replacement | 29 (12.9) | 3 (6.8) | 0 (0.0) |
| MV Repair | 196 (87.1) | 41(93.2) | 0 (0.0) |
| **Aortic Valve (AV) Dysfunction** |  |  |  |
| AV Insufficiency | 19 (8.4) | 4 (9.1) | 1 (5.0) |
| AV Stenosis | 10 (4.4) | 2 (4.5) | 0 (0.0) |
| AV Surgical Intervention | 10 (4.4) | 1 (2.3) | 0 (0.0) |
| **Coronary Artery Disease** | 79 (35.1) | 14 (31.8) | 3 (15.0) |
| Intervention: CABG | 24 (10.7) | 4 (9.1) | 1 (5.0) |
| Intervention: Stent | 11 (4.9) | 2 (4.5) | 0 (0.0) |
| **Atrial Fibrillation** | 52 (23.1) | 12 (27.3) | 5 (25.0) |
| Intervention: Ablation | 29 (12.9) | 6 (13.6) | 0 (0.0) |
| **New York Heart Association Class III-IV** | 93 (41.3) | 16 (36.4) | 9 (45.0) |
| **Comorbidities** |  |  |  |
| Hypertension | 127 (56.4) | 27 (61.4) | 9 (45.0) |
| Hyperlipidemia | 92 (40.9) | 20 (45.5) | 8 (40.0) |
| Diabetes Mellitus | 18 (8.0) | 3 (6.8) | 5 (25.0) |
| Renal Insufficiency | 16 (7.1) | 3 (6.8) | 6 (30.0) |
| **Medications** |  |  |  |
| Aspirin | 91 (40.4) | 14 (31.8) | 8 (40.0) |
| Beta Blockers | 94 (41.8) | 20 (45.5) | 9 (45.0) |
| ACE Inhibitors | 69 (30.7) | 11 (25.0) | 8 (40.0) |
| Statin | 85 (37.8) | 20 (45.5) | 7 (35.0) |
| Diuretic | 69 (30.7) | 14 (31.8) | 11 (55.0) |
| Plavix | 2 (0.9) | 1 (2.3) | 1 (5.0) |
| Warfarin | 26 (11.6) | 6 (13.6) | 9 (45.0) |
| Other Anticoagulant | 23 (10.2) | 5 (11.4) | 1 (5.0) |
| ARB | 33 (14.7) | 6 (13.6) | 2 (10.0) |
| Calcium Channel Blocker | 33 (14.7) | 5 (11.4) | 1 (5.0) |
| Anti-Arrhythmic | 16 (17.1) | 4 (9.1) | 2 (10.0) |
| SSRI | 20 (8.9) | 5 (11.4) | 4 (20.0) |
| Other Psychiatric | 17 (7.6) | 4 (9.1) | 2 (10.0) |

### The Optum Database—de-identified patient data for multivariate analyses (Table S2)

OptumInsights Informatic Data Mart (Optum) is a large private payer administrative claims database that comprises a geographically diverse population and contains approximately 12-15 million patients within the United States. This database was accessed through a licensed site at the Leonard Davis Institute, University of Pennsylvania. 9,441 patients over 18 years of age with mitral valve regurgitation who underwent surgery and were included in the Optum Database from 2004 to 2016, were studied. We found 6,751 (71.5%) patients that underwent mitral valve repair and 2,690 (28.49%) patients that underwent mitral valve replacement. Patients with mitral valve stenosis prior mitral valve surgery, and endocarditis were excluded. Additionally, we excluded any patient that underwent a concomitant coronary artery bypass graft (CABG) procedure at the time of the mitral valve surgery in order to exclude patients with either possible ischemic or functional mitral valve regurgitation. This was done in an attempt to estimate the extent of degenerative mitral regurgitation, because the specific types of mitral regurgitation are not coded in the Optum Database. Age at surgery is a whole number in the Optum Database, as birthdates are not entered in order to protect patient identity. SSRI use was ascertained as described above. Aspirin use prior to surgery, since this drug is not prescribed, is not recorded in the Optum dataset.

### RNA isolation from mitral valve leaflets and qRT-PCR methodology

Following surgical pathology evaluation and sampling for routine histology examination, available samples of MR leaflet specimens and normal mitral leaflets were immediately flash frozen in liquid nitrogen. RNA samples suitable for microarray and qRT-PCR analyses were isolated from MR surgical specimens using the RNeasy Fibrous Tissue Mini Kit (QIagen, Germantown, MD). RNA concentrations were measured by Nanodrop technology (Thermofisher), and RNA integrity was assessed with the Agilent 2100 Bioanalyzer (Agilent, Santa Clara, CA). cDNA was prepared from RNA using the RT2 first strand system (Qiagen).

qRT-PCR analyses were performed using RT2 Profiler Arrays (Qiagen) with specificity for either Dopamine-5HT related genes or TGFβ related gene expression. Each array contained a panel of proprietary controls to monitor genomic DNA contamination as well as the first strand synthesis, and real-time PCR efficiency (Qiagen). The qRT-PCR data were used to analyze and compare MR and normal mitral leaflet gene expression patterns. The Dopamine-5HT RT2 profiler array contained 96 target genes, as well as housekeeping genes for normalization that included: Actin-beta, Beta-2-microglobulin, Glyceraldehyde-3-phosphate dehydrogenase, and Ribosomal protein-large-P0. The TGFβ RT2 Profiler Array contained 96 target genes, with data normalization utilizing the following housekeeping genes: Actin-beta, Beta-2-microglobulin, Glyceraldehyde-3-phosphate dehydrogenase, and Ribosomal protein-large-P0. Gene expression levels were computed by normalizing with the housekeeping genes using the delta cycle threshold (ΔCt) methodology, where ΔCt means the Ct of the target gene minus the average Ct of the house keeping genes described above. Fold changes were calculated using ΔΔCt method, 2^−ΔCt for MR leaflets^/2^−ΔCt for normal leaflets^, thereby calculating fold regulation comparing MR to normal mitral leaflet gene expression.

### Mitral valve interstitial cell studies

Primary mitral valve interstitial cell cultures were established for cells derived from excised mitral leaflets sub-populations (Tables S7 and S8) of the clinical study groups (Methods S1) for both MR leaflets, and normal leaflets using established methodology ^26^. qRT-PCR studies of these cultures compared gene expression levels of SLC6A4 and HTR2B with and without the addition of Fluoxetine to the cultures. These qRT-PCR studies also included measurements of α-smooth muscle actin and vimentin, as activation markers, in RNA from both the MVIC post culture and the valve leaflets that were the sources of the MVIC (Methods S1).

### Platelet studies (Methods S1)

The effects of Fluoxetine and Aspirin on ADP induced platelet activation were studied on fresh blood samples obtained per Children’s Hospital of Philadelphia IRB Protocol 12-008608 from healthy human subjects (Tables S9 and S10) with no history of heart valve disease and on no medications (Methods S1). Platelet studies utilized a Chandler Loop apparatus ^32, 33^ (Methods S1). Endpoints per flow cytometry were total platelet count as an index of aggregation for each ADP dose, and P-Selectin. Plasma 5HT levels (Methods S1) for each ADP dose, treated and untreated were measured using mass spectroscopy with isotope dilution methodology ^34^.

### Statistical methods

Data were expressed as means ± standard deviations. Clinical data, both from the primary patient population and the Optum dataset, were assessed using Cox’s proportional hazards model where the response variable was the age at mitral valve surgery. Thus, the MVA approach used needs clarification, since cross sectional data were used to determine age at surgery rather than the ideal endpoint of risk of MR. However, since MR surgery is only performed for end stage disease, this approach was empirically used with the caveat just stated. The rationale for this is that cardiac surgery for MR is the only treatment option, and because of the significant risks involved, it is deferred as long as possible. MR surgery is never elective. Younger age at surgery indicates more rapidly progressive disease. The covariates were then pared down to a subset that were of statistical significance. qRT-PCR data were subjected to analyses based on ΔCt methodology, as described above, and Student’s t tests for significance per the manufacturer’s analytical software (Qiagen), with results plotted as fold change showing 95% confidence limits. Paired t tests were used for the platelet studies. Cell culture qRT-PCR data were analyzed using Kruskal Wallis methodology. All reported p values were calculated on the basis of two-sided tests, and a p value of less than 0.05 was considered to indicate statistical significance.

## Results

### The degenerative mitral regurgitation study population

225 patients requiring cardiac surgery for MR were studied (Table 1). The age range at the time of surgery was 24 to 89 years, and 71.1% of the patients were men with a mean age at surgery of 62.9 years; female patients were not significantly older with a mean age of 63.8 years. The majority of patients underwent mitral valve repair, 87.1%, and the remainder had mitral valve replacements with prostheses (Table 1). Overall 41.3% of these patients were determined to be New York Heart Association (NYHA) Class III or IV. Other frequent cardiac comorbidities (Table 1) included associated coronary artery disease, not related to MR (35.1%), aortic valve disease (12.8%), and atrial arrhythmias (23.1%). Hypertension and hyperlipidemia were also highly prevalent (Table 1).

### Multivariate analyses according to age at the time of surgery for degenerative mitral regurgitation (Figure 1)

#### The primary 225 patient population results

Cox model analyses (Figure 1A, and Table S1) revealed that a number of comorbidities reduced the hazard of MR surgery; in other words, these comorbidities were associated with surgery at an older age. These comorbidities include coronary disease (p<0.001), concomitant aortic valve disease (p<0.001), heart failure (p=0.01), atrial fibrillation (p=0.02), renal insufficiency (p<0.001), and hyperlipidemia (p=0.0088). Male gender was significantly associated with surgery at a younger age (Figure 1A), and there was a male predominance in the population overall (Table 1). It is notable that all of the variables in Figure 1A were significant in accounting for the number of original variables that were considered in terms of the false discovery rate. Aspirin therapy was administered to more than 40% of the population (Table 1 and Figure 1A), and after adjusting for co-variates, was associated with a reduction in the hazard of having MR surgery at a younger age (p=0.0055). SSRI administration prior to MR surgery was documented in 20 of the 225 patients (Table 1), and after adjusting for co-variates in the Cox model, was significantly associated with increasing the hazard of having MR surgery at a younger age (Figure 1A, p=0.0183).

**Figure 1:**
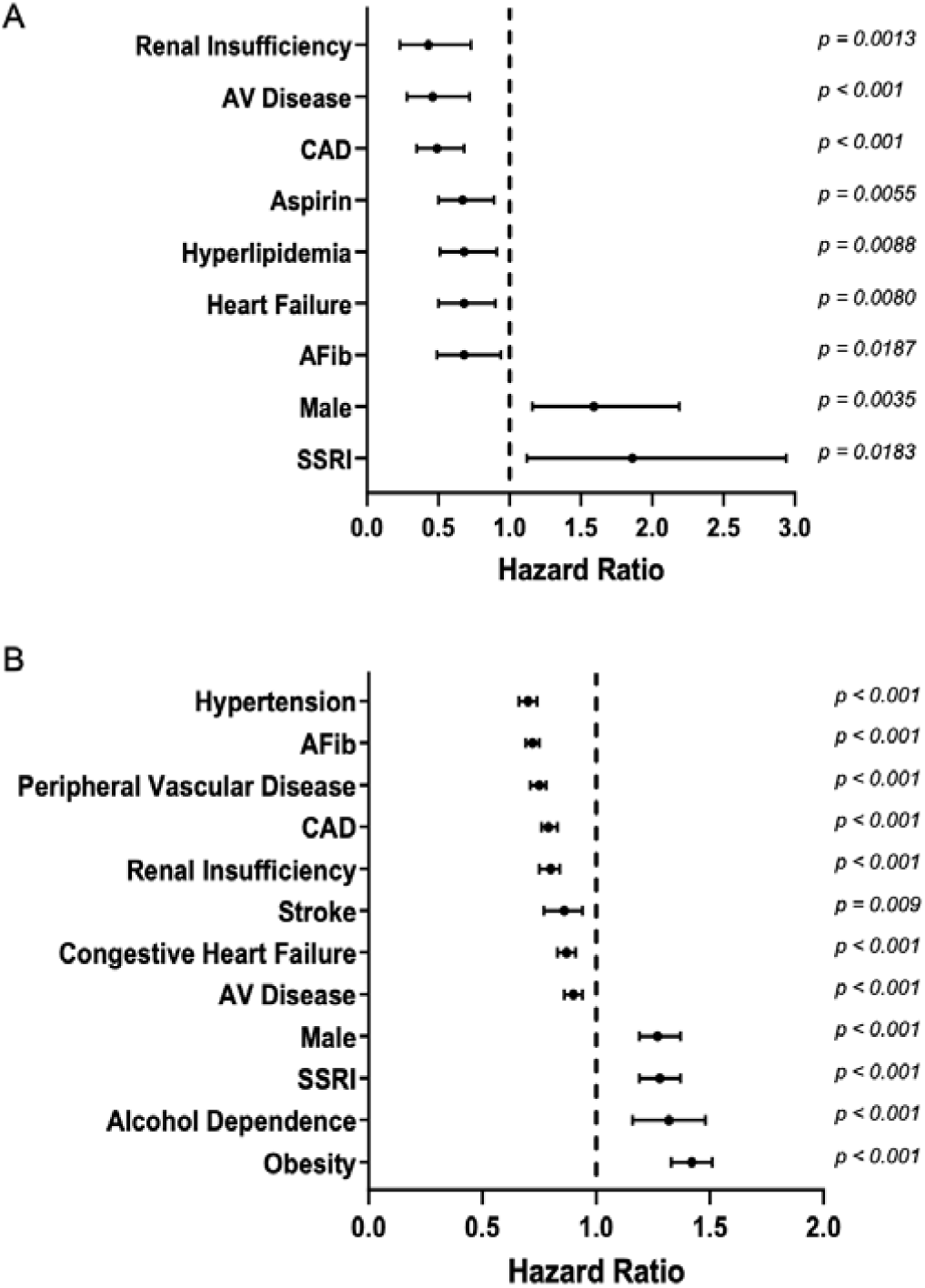
Multivariate Cox model analyses results of patients with degenerative mitral regurgitation requiring cardiac surgery. Thus, data were assessed using statistical methodology that employed Cox’s proportional hazards model where the response variable was the age at mitral valve surgery. Co-variates are shown with 95% confidence limits reflecting the hazard of requiring surgery at younger ages (hazard ratio>1.0) compared to older age at surgery (hazard ratio<1.0). Shown are Forest plots with p values for: A) The primary 225 patient population, and B) The Optum 9441 patient population, that includes patients not specifically coded for degenerative mitral regurgitation (see Methods). Abbreviations used are: AV, aortic valve disease; CAD, coronary artery disease; AFib, atrial fibrillation; SSRI, serotonin reuptake inhibitor.

#### Optum Database results (Figure 1B)

This dataset included a number of co-variates for analyses that were present in the primary 225 patient population and others that were not mutually available (Figure 1B, Table S2). In the Optum population (9,441 patients with mitral valve regurgitation undergoing surgery, Table S2), SSRI use (Figure 1B) was also significantly associated with increasing the hazard of having MR surgery at a younger age (p<0.001). In addition, the Optum results showed that concomitant aortic valve disease, congestive heart failure, atrial fibrillation, coronary artery disease (without concomitant bypass surgery), nd renal insufficiency, were also significantly associated (p<0.001) with a hazard ratio favoring MR surgery at an older age (Figure 1B, Table S2). Obesity and alcohol dependence in the Optum data (Figure 1B), co-variates not available in the primary patient dataset (Figure 1A), were significantly associated (p<0.001) with a hazard ratio favoring surgery at a younger age (Figure 1B, Table S2). Male gender was significantly associated with surgery at a younger age (Figure 1B), in agreement with the results in the primary surgery population (Figure 1A); 56.7% of the MR subjects in the Optum dataset were males (Table S2). In the Optum population hypertension and peripheral vascular disease were each significantly associated (p<0.001) with a hazard for surgery at an older age (Figure 1B, Table S2).

### Differential gene expression in degenerative mitral regurgitation leaflets compared to anatomically normal mitral leaflets

#### 5HT related gene expression results (Figure 2)

MR leaflets were compared to normal mitral leaflets by qRT-PCR analyses (Figure 2 and Tables S3-S6). RNA samples of suitable quality for qRT-PCR analyses were available from 44 MR leaflet samples of the 225 MR cases. The demographics of this 44 patient MR data set (Table 1) were comparable to the overall population (Table 1). There were 31 male and 13 female patients in the subpopulation for PCR analyses. Of the 44 MR cases, 36 involved resections of the posterior mitral leaflets, and the remainder were anterior leaflet repairs. It should be noted that this subgroup included 14 patients (31.8%) with coronary disease (Table 1), who were clinically assessed as not having MR related to this co-morbidity. The 20 anatomically normal mitral leaflets obtained from transplant cases included 13 men and 7 women (Table 1). This group (Table 1) included 3 subjects (15.0%) with coronary disease, and none with mitral regurgitation.

**Figure 2:**
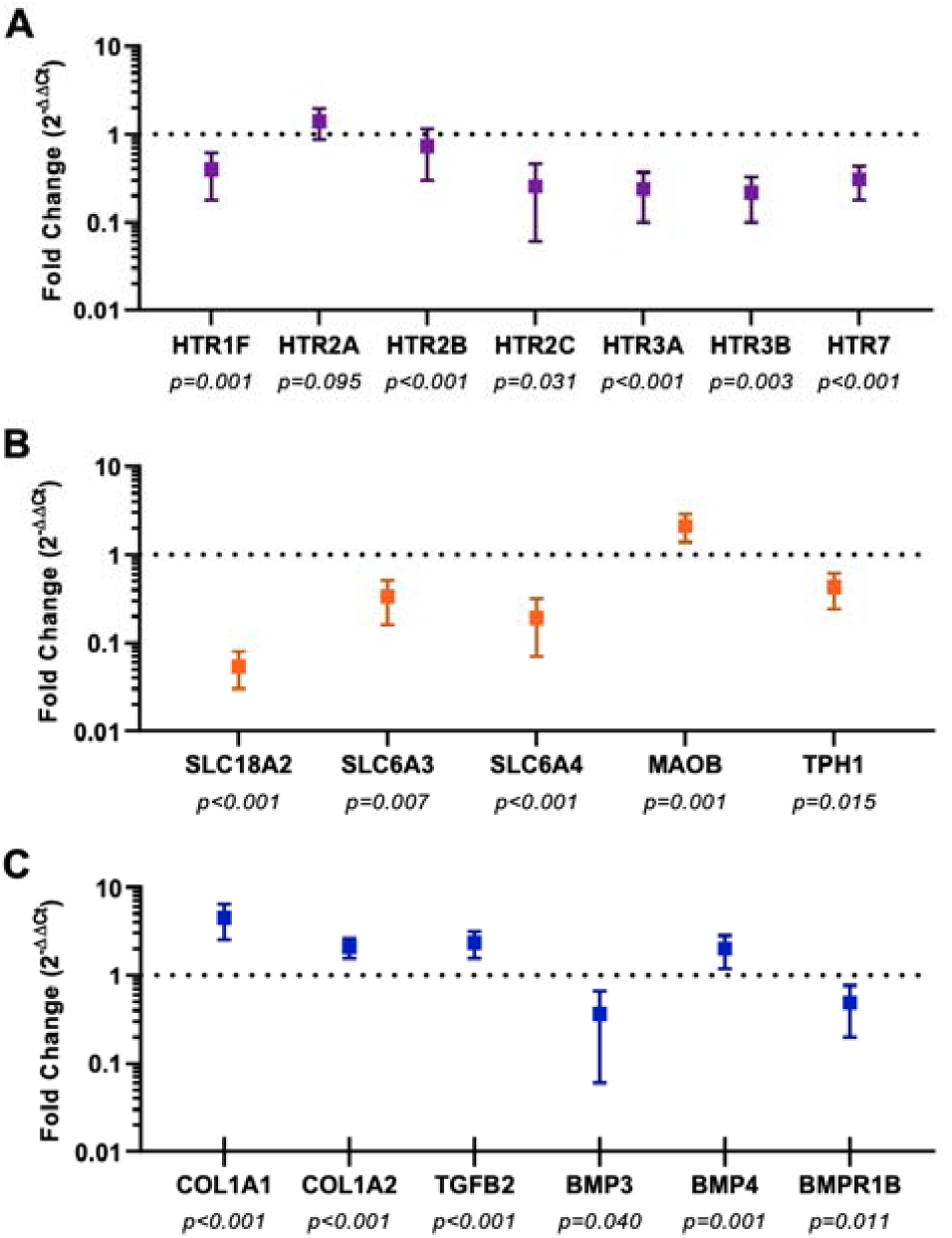
Quantitative reverse transcriptase polymerase chain reaction (qRT-PCR) results shown as fold regulation changes in degenerative mitral regurgitation (MR) leaflets versus anatomically normal leaflets (see Tables S5 and S6 for complete data and gene symbol descriptions): A) Serotonin receptor (HTR) expression level comparisons MR versus normal leaflet results; B) Serotonin transporter (SLC6A4) and other monoamine transporter expression levels comparing MR to normal leaflets; C) Serotonin synthetic genes (tryptophan hydroxylase, TPH) and monoamine oxidase expression (MOAB) fold regulation, MR compared to normal; D) TGFβ superfamily genes and collagen expression in MR compared to normal mitral leaflets. qRT-PCR data were subjected to statistical analyses based on ΔCt (Methods) and Student’s t tests (two sided) were used for assessing significance. Results a re shown as fold changes with p values. N.S. indicates a p value that is not significant (p>0.05).

5HT-related qRT-PCR results (Figures 2A, B, and C; Table S3) demonstrated a significant down regulation of 5HT receptors HTR1F, HTR2B, HTR2C, HTR3A, HTR3B and HTR7 (Figure 2A). However, HTR2A was not significantly different in MR compared to normal, and HTR2B, while significantly down regulated (Figure 2A), demonstrated 1.37 fold down regulation. In addition there was significant down regulation of monoamine transporter genes (Figure 2B, Table S3) SLC6A4 (−5.22 fold, p<0.001), vesicular monoamine transporter 2 (SLC18A2, −18.57 fold, p<0.001), and the dopamine transporter (SLC6A3, −3.00 fold, p=0.007). Serotonin synthetic genes (tryptophan hydroxylase, TPH1 and TPH2 (Figure 2C) were significantly down regulated in MR, and monoamine oxidase-B expression (MOAB) was significantly upregulated, MR compared to normal (Figure 2C, and Table S4).

#### TGFβ related gene expression results (Figure 2D)

qRT-PCR data (Figure 2D, and Table S4) demonstrated significant upregulation of a number of genes in MR compared to normal. Genes upregulated in MR (Figure 2D, Table S6) included collagens-1 alpha-1&2 (COLA1A1, and COL1A2, p<0.001) and TGFβ2 (TGFB2, p<0.001). Down regulation (Figure 2D, Table S4) was noted for the following genes of interest (Figure 2D, Table S4): Bone morphogenic protein 3 (BMP3, p=0.040), and BMP receptor 1B (BMPR1B, p=0.011). BMP4 was upregulated 2.02 fold in MR (p=0.001). It was also observed that TGFβ1 and the TGFβ-receptor-1 were not differentially expressed comparing MR and normal leaflets (data not shown).

#### Cell Culture Studies (Figure 3)

Cell culture studies of human mitral valve interstitial cells (MVIC) investigated SSRI related effects on both SLC6A4 expression. These studies were based on prior research concerned with the central nervous system that demonstrated down regulation of SLC6A4 due to Fluoxetine, a SSRI ^35, 36^ in both zebrafish ^35^ and rats ^36^. In addition, HTR2B expression was an endpoint because of previous studies demonstrating the potential importance of this receptor in valve disease ^7, 11, 26, 37^. Primary human mitral valve interstitial cell (MVIC) studies, using cells cultivated from leaflets harvested from both MR cases (Table S5) and normal leaflets (Table S6), were carried out under full serum conditions comparing each cell line with or without exposure to Fluoxetine (10μM). Endpoints were changes in HTR2B and SLC6A4 expression levels measured with qRT-PCR. α-smooth muscle actin and vimentin expression levels were used as activation markers (Methods S1). There were no significant differences in activation markers’ expression between diseased and normal leaflets that provided MVIC for these studies (data not shown). However, Fluoxetine resulted in a significant down regulation in α-smooth muscle actin in MVIC from normal leaflets, but not MR derived MVIC (Figure 3). Vimentin was significantly upregulated in MVIC by Fluoxetine in MR derived cells, but was not significantly affected in MVIC from normal leaflets (Figure 3). 5HT related results showed that Fluoxetine caused a significant down regulation of SLC6A4 in both MR (p=0.001) and normal (p=0.002) MVIC (Figure 3). HR2B demonstrated the greatest increase in expression levels, 174.3% in MR MVIC with Fluoxetine exposure (p=0.005); a modest, 112%, but significant increase in HTR2B expression was observed with Fluoxetine in MVIC from anatomically normal leaflets (Figure 3, p=0.01).

**Figure 3:**
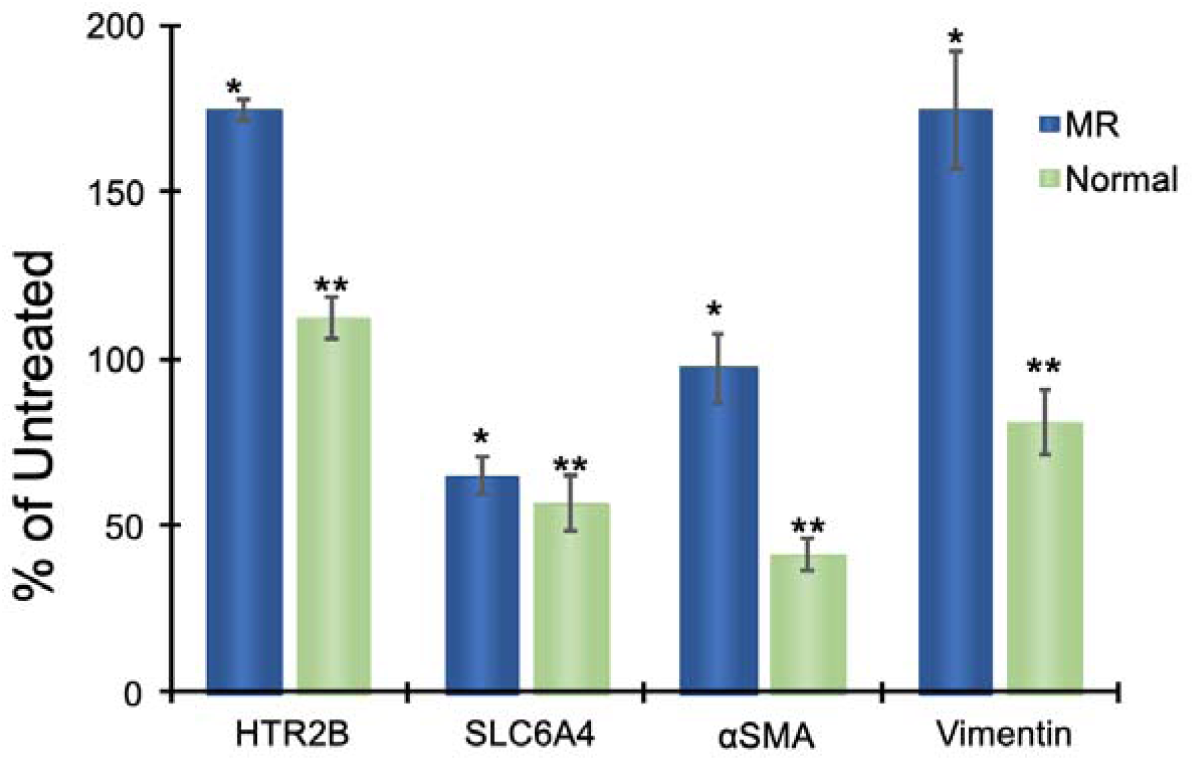
Changes in human mitral valve interstitial cell gene expression levels with Fluoxetine administration (1µM): qRT-PCR results comparing 5HT receptor 2B (HTR2B) and 5HT-transporter (SLC6A4); activation marker data are also displayed for alpha-smooth muscle actin (αSMA) and vimentin. Data shown indicate percentage change in expression levels comparing Fluoxetine treated cultures from the same human subject to untreated: HTR2B, *p=0.005, **p=0.001; SLC6A4, *p=0.001, **p=0.002; αSMA, *p=>0.05, **0.006; vimentin, *p=0.04, **p=>0.05. Results shown are means ± standard deviations.

#### Platelet Results (Figure 4)

Platelet studies used an ex vivo blood flow system, the Chandler Loop apparatus ^32, 33^, to study ADP induced platelet activation as a model of MR associated platelet activation ^27, 38^. In these platelet experiments the effects of Fluoxetine, a SSRI, and separately, Aspirin, an inhibitor of platelet activation, on platelet number, activation and 5HT release were assessed. Fresh human blood samples were studied (Tables S7 and S9) that were anticoagulated with citrate ^33^. Fluoxetine (10μM) and Aspirin (250 μM) were studied separately in samples from healthy human subjects, five subjects in each drug study group; these individuals had no known diseases and were receiving no medications. The procedure followed was to compare a non-treated blood sample to a drug treated sample from the same subject. Following the addition of the medication to the blood sample, four hours duration simultaneous runs were performed for both treated and untreated samples from each subject at 37C in the Chandler Loop. This was followed by ADP induced platelet activation of samples removed from the Chandler Loop, as a model of MR associated platelet activation ^27, 38^, over a dose range of ADP (Figure 4). Fluoxetine compared to untreated resulted in significant enhancement of ADP-induced platelet activation, demonstrating increased P-Selectin levels, reduction in platelet counts, indicating aggregation, and increased plasma 5HT levels, both prior to ADP and as a result of ADP administration (Figures 4A-C). Aspirin administration using the same protocol was observed to cause an inhibition of ADP-induced platelet activation as shown with reduced P-Selectin levels, and higher platelet counts compared to untreated (Figures 4D, E). In the presence of Aspirin, 5HT levels did not significantly increase with ADP induced activation (Figures 4F).

**Figure 4:**
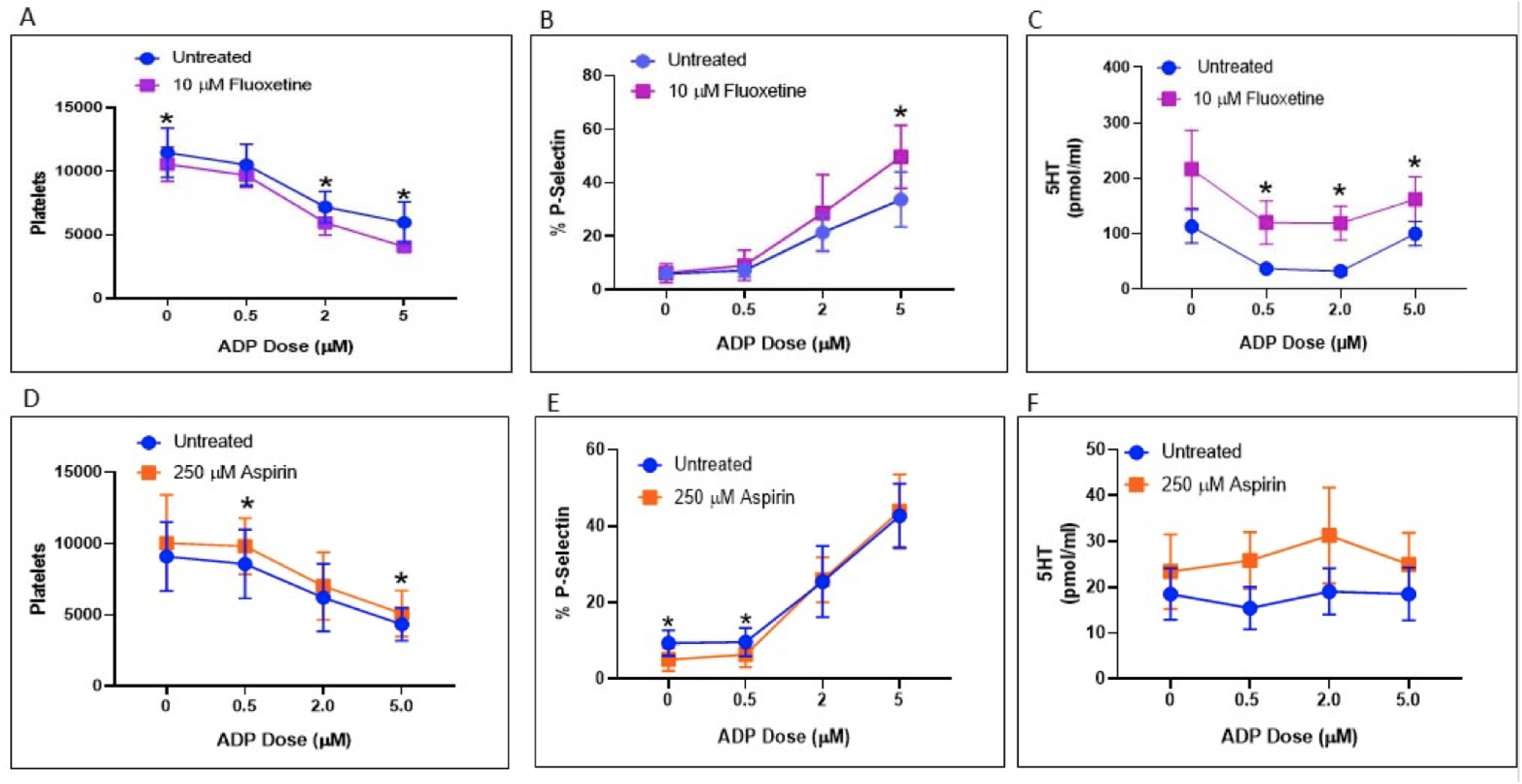
Platelet studies investigating the effects of Fluoxetine and Aspirin. Fluoxetine (10µM) was added to blood samples from 5 healthy donors (Fig. 3A-C), and compared to untreated blood samples from each of the individual donors. Chandler Loop runs of 4 hours of these samples, treated and untreated, were followed by removing samples from the loop, and the addition of ADP to induce platelet activation using the dosages shown. A) Effects of Fluoxetine on total platelet count, where, *0.0 µM ADP, p= 0.046, *2.0 µM, p=0.042, *5.0 µM, p=0.041; B) Percentage of platelets that were P-selectin positive comparing Fluoxetine to untreated, where *5.0 µM ADP, p= 0.044; C) Serotonin (5HT) levels comparing Fluoxetine to untreated where *0.5 µM ADP, p=0.030, *2.0 µM ADP, p= 0.041 and *5.0 µM ADP, p= 0.050. Aspirin studies (Fig. 3D-F) used the same methodology as the Fluoxetine experiments, with 5 subjects, but with Aspirin added at a concentration of 250 µM and compared for each of the individual subjects to untreated blood. D) Effects of Aspirin versus untreated on platelet counts where *0.5 µM ADP, p=0.0029, *5.0 µM ADP, p= 0.020; E) the percentage of cells that were P-selectin positive comparing Aspirin to untreated, where 0.00 µM ADP, p=0.0031, *0.5 µM ADP, p=0.024 F) 5HT l vels comparing Aspirin to untreated demonstrated no significant differences between Aspirin treated and untreated. Data shown are means with standard errors as error bars. Statistical methodology used paired t tests for these platelet studies. Paired t-tests, two sided, compared each subjects blood sample, treated and untreated; p values were calculated accordingly.

## Discussion

The present results indicate that the progression of MR is influenced by both diminished SLC6A4 expression and SSRI administration (Figures 1 and 2). This was demonstrated by the multivariate analyses of clinical data, indicating a significant SSRI effect, with an increased hazard for surgery at a younger age in patients taking SSRI (Figure 1), and the major differences in 5HT gene expression patterns in MR leaflets compared to normal, with significant down regulation of SLC6A4 and other monoamine transporters (Figure 2), that would result diminished 5HT deactivation thereby enhancing 5HT activity. 5HT receptor signaling by HTR2A and HTR2B, which are not down regulated in MR (Figure 2) upregulates TGFβ1^12, 39^, resulting in increased extracellular matrix (ECM) protein production^12, 39^ that contributes to the progression of MR.

The TGFβ related results in our qRT-PCR data (Figure 2C) demonstrate increased ECM and TGFβ related gene expression pattern changes that support the hypothesis that increased HTR signaling with downstream effects has occurred over time in the MR leaflets. Thus, relatively increased 5HT due to either reduced SLC6A4 or pharmacologic inhibition would enhance HTR signaling, as has been shown experimentally ^21^ (Figure 3). In addition, Filamin-A, an important cytoskeletal protein involved with organization of ECM, could also be affected by the mechanisms of interest in the present studies. Filamin A mutations cause an X-linked myxomatous mitral valve disease ^31^, and mice with an endocardial specific conditional Filamin-A deletion demonstrate thickening of mitral leaflets^40^. Diminished 5HT internalization due to reduced SLC6A4 activity results in diminished serotonylation of Filamin A in the mouse model ^40^, thereby impairing Filamin-A structural interactions with actin, enhancing ECM pathologic remodeling in mitral leaflets. A comparable mechanism could be affecting MR leaflets due to reduced monoamine transporter capacity (Figure 2B).

HTR2B, that is not significantly reduced in MR leaflets compared to normal mitral leaflets (Figure 2) is thought to be of key importance in 5HT related valvulopathies ^6^, and is also of interest mechanistically in fibrotic diseases ^13^. HTR2B signaling upregulates TGFβ1 in heart valve interstitial cells as well, resulting in increased ECM ^12, 39, 41^. In the present studies MVIC experiments demonstrated both significant upregulation of HTR2B and down regulation of SLC6A4 in response to Fluoxetine incubations (Figure 3). Comparable SSRI mediated down regulation of SLC6A4 has been observed in vivo in central nervous system studies in both zebrafish ^35^ and rats ^36^. SSRI upregulation of HTR2B has not been previously reported. These changes in gene expression patterns in the present MR studies would hypothetically enhance 5HT interactions with increased HTR2B signaling capacity. MVIC are heterogeneous and have been incompletely characterized. The MVIC phenotype of most importance in HTR2B signaling at present is unknown. However, there is compelling evidence from clinical pathology analyses ^37, 42^ and mouse model studies ^37, 43^ that bone marrow derived endothelial progenitor cells expressing HTR2B are present in both normal and regurgitant mitral leaflets, and are mobilized from bone marrow through HTR2B signaling ^37^. These bone marrow derived MVIC may contribute substantially to HTR2B signaling involved in the pathophysiology of MR.

The importance of platelets in 5HT mechanisms related to heart valve disease has been explored to a limited extent ^27, 28, 30, 38^. Platelets are the primary carriers of 5HT in blood. 5HT produced by chromaffin cells in the small intestine is taken up by platelets in the capillary circulation through the coordinated action of platelet SLC6A4, and SLC18A2, packaging 5HT in platelet dense granules ^29^. Platelet activation results in the release of 5HT from dense granules ^29^. 5HT release due to platelet activation has several significant effects including local vasoconstriction and enhanced platelet aggregation through signaling via platelet HTR2A and HTR2B ^44^. Although MR has been shown to result in a chronic state of platelet activation ^27, 28^, 5HT levels have not been studied in MR patients. However, aortic stenosis also results in enhanced platelet activation, and increased 5HT levels have been documented in a clinical study comparing aortic stenosis patients to those without valvular heart disease ^30^. In our platelet studies, SSRI administration resulted in enhanced ADP induced platelet activation accompanied by increased 5HT levels (Figure 4). Anti-platelet agents such as Aspirin (Figures 1 and 4) by inhibiting platelet activation, that is also associated with both cytokine release from alpha granules, and 5HT from dense granules ^45^, could hypothetically have an impact on the progression of MR by mitigating these platelet related contributions to the pathophysiology.

The present studies have several limitations inherent in a study design driven by selecting patients for investigation who require cardiac surgery. This may have resulted in a population of MR patients with very different 5HT susceptibilities and gene expression patterns in their mitral leaflets than less severely ill subjects. In addition, although medication history was accurately ascertained, it was not possible to obtain comprehensive information about duration of use or dosing. Furthermore, as noted in Methods, since the Optum data set does not distinguish between degenerative mitral regurgitation, and mitral regurgitation due to ischemic heart disease, patients requiring concomitant coronary bypass surgery were excluded to hypothetically adjust for this. However, 49.0% of the Optum patients analyzed with MR (Table S2) had coronary disease, versus 35.1% (Table S1) in the 225 patient population. Nevertheless, both MVAs (Figure 1) showed that coronary artery disease was associated with mitral valve surgery at an older age, in contrast to the SSRI effect associated with MR surgery at a significantly younger age (Figure 1). Also, Aspirin use was not reported in the Optum dataset, since this agent is not a prescribed drug, and therefore has no insurance costs. Thus, there is no opportunity to confirm the MVA results (Figure 1A) for Aspirin in the Optum data. The qRT-PCR studies while highly significant were carried out on samples obtained from end stage MR valve leaflets, in comparison to normal heart valve leaflets that were obtained from patients in heart failure requiring heart transplants. In addition 4 of the normal leaflets in the qRT-PCR data set were from patients taking SSRI. Their qRT-PCR results did not differ from the rest of the population (data not shown); however, this small sample size limits any conclusions about an SSRI effect. These limitations cannot be overcome in a human population. For qRT-PCR leaflet samples, RNA was only available on 44 of the 225 MR patients, and this limitation relates to a clinical reality of often confining surgery to annuloplasty, conserving valve tissue in many cases, sampling required for surgical pathology review, and deterioration of RNA prior to processing. Our platelet studies were carried out on healthy subjects on no medications, rather than investigating a MR population. This was necessary to focus on specific 5HT platelet mechanisms of interest and avoid for now the complex comorbidities (Table 1) associated with a MR population. Platelet release of 5HT with Fluoxetine administration and associated platelet activation have been observed in studies of patients with depression ^46^, and these results are in agreement with our data.

It is concluded that SLC6A4 contributes to the pathophysiology of MR (Figure 5) through enhanced HTR signaling in MVIC in MR leaflets, due to relatively greater 5HT exposure arising from diminished SLC6A4, direct inhibition of SLC6A4 by SSRI, and reduced SLC18A2 expression. SSRI can further enhance 5HT effects in MR through upregulation of HTR2B and down regulation of SLC6A4 in MVIC. Increased platelet derived 5HT resulting from SSRI enhances platelet activation. Inhibiting 5HT effects through the use of agents, such as Aspirin, that reduce the level of platelet activation and the related release of 5HT represents a potential therapeutic direction.

**Figure 5:**
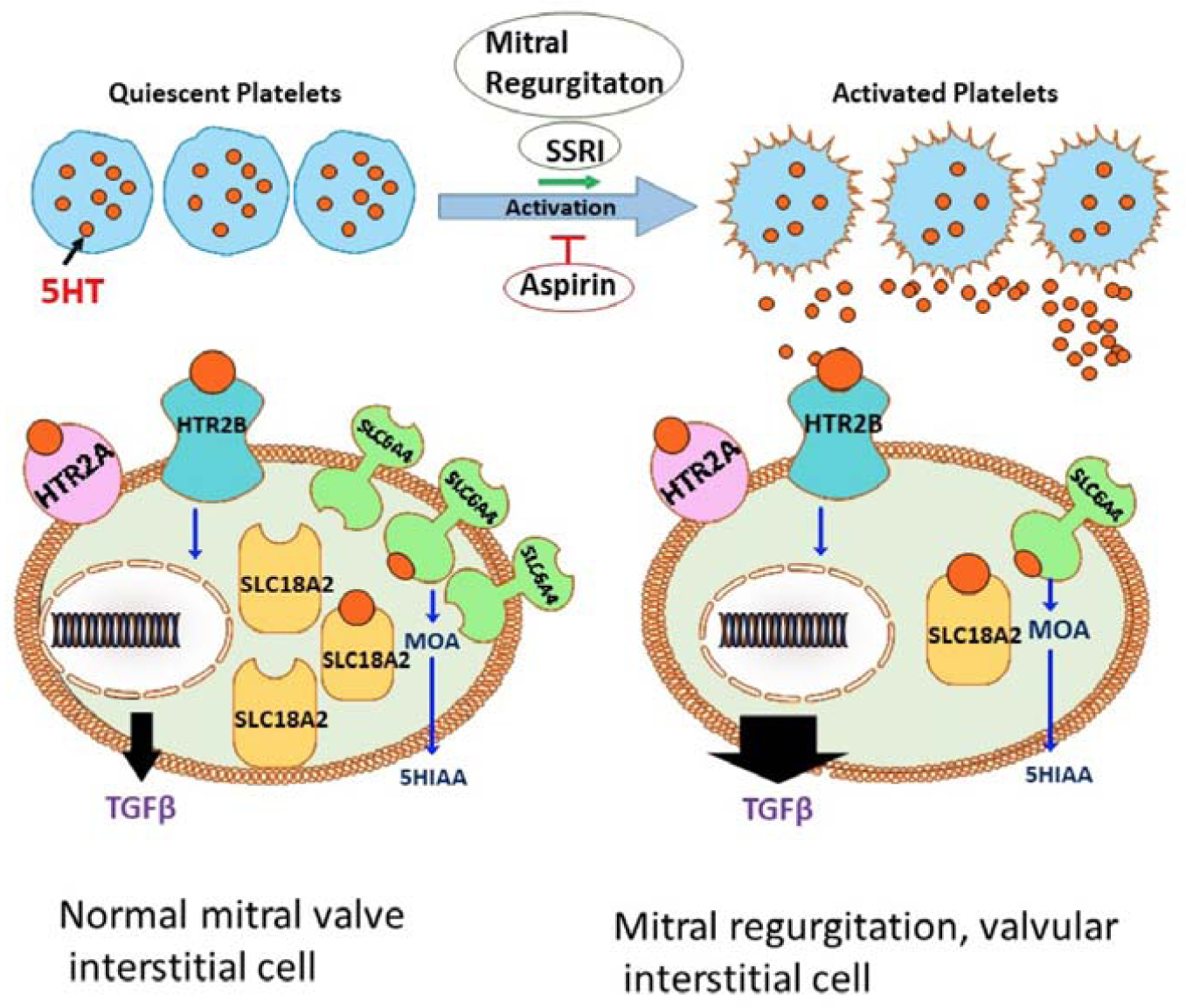
Serotonin transporter (SLC6A4) mechanisms that influence the mitral valve interstitial cell (MVIC) response in the pathogenesis of degenerative mitral regurgitation (MR): Disease progression is associated with diminished expression of both SLC6A4 expression (Figure 2) and vesicular monoamine transporter-2 (SLC18A2, see Figure 2). This results in diminished deactivation of 5HT by monoamine oxidase (MOA) that would otherwise reduce 5HT levels producing 5-hydroxyindolacetic acid (5HIAA). Platelet activation due to MR^27^also serves to elevate 5HT (Figure 3). SSRI administration, by both blocking 5HT uptake by MVIC^21^ increases HTR signaling, and related upregulation of TGFβ mechanisms via receptors HTR2A ^12, 39^ and HTR2B ^11^, that are not down regulated in MR (Figure 2). SSRI can also down regulate SLC6A4 expression (Figure 3), while upregulating HTR2B in MVIC (Figure 3) further enhancing 5HT effects. Fluoxetine, a representative SSRI, enhances platelet activation, and increases platelet secretion of 5HT thus enhancing HTR signaling (Figure 3A-C). Aspirin administration inhibits platelet activation (Figure 3) and thus, could mitigate MR progression by reducing platelet activation associated 5HT release from platelets (Figure 3D-F), as well as cytokine release, including TGFβ, from the alpha-granules of activated platelets. ^47^

## Author contributions

Conception and planning of experiments: RJL, GF; funding secured by RJL and GF; writing by: RJL, EGF, EC, HSS, SJS, CRB, AMK, GF; close manuscript review: RJL, AEA, JGG, SJK, RCG, LR, SJS, GF; experiments performed by: RJL, EGF, EC, HJS, VVI, ES, SJK, IN, CRB, AMK, GF; explants, non-commercial material and supplies provided by: AEA, JGG, NR, SJK, RCG; data and statistical analysis: RJL, EGF, EC, HJS, SJK, IN, CRB, AMK, GF; interpretation of results: RJL, EGF, EC, HJS, SJK, CRB, GF; preliminary studies performed by: RJL, EC, HJS, VVI, AEA, ES, SJK, IN, SKS, GF. All authors provided critical feedback and helped shape the research, analysis and writing the manuscript.

The authors thank Ms Susan Kerns for her assistance in the submission of the manuscript. The authors also thank Peter W. Groeneveld, MD, University of Pennsylvania, for his helpful advice about planning the national database studies. The authors are grateful to Kenneth Margulies, MD and Kenneth Bedi, The University of Pennsylvania, for their assistance in obtaining anatomically normal mitral valve leaflets as permitted by IRB Protocol #802781. We thank Y. Daikhin, O. Horyn and Ilanna Nissim for performing the 5HT measurements in the Metabolomics Core Facility, The Children’s Hospital of Philadelphia.

## Methods S1

### Cell Culture Methodology

Human mitral valve (MV) interstitial cells (MVIC) were isolated from MV leaflet tissue explanted during MR surgery (mitral regurgitation MVIC) or heart transplant (normal leaflet MVIC). For human subjects demographics see Tables S7 and S8. Leaflets were minced and digested with 1 mg/ml collagenase type 2 (Worthington Biochemical Corporation, Lakewood, NJ) and 100 IU/mL hyaluronidase (Worthington) in complete growth medium for 16h-24h at 37°C. After digestion, the cell suspension was pelleted in a centrifuge (5 min, 2000 rpm). Cells were cultivated in complete growth medium (Advanced DMEM containing 4.5g/ml glucose, supplemented with 10% FBS, 4 mM L-glutamine, 1% penicillin/streptomycin, and 1μg/mL amphotericin-B. Phenotypic validation of mitral regurgitation and normal MVICs was performed by assessing expression levels of MVIC markers using qRT-PCR (α-smooth muscle actin Hs00426835_g1, and desmin, Hs00157258_m1, both FAM-based Taqman probes from Thermo) and confirming the absence of expression of the endothelial cell marker CD31 (Hs01065279_m1, Thermo). Cryorecovered MVICs at passages 2-4 were used for all experiments. MVICs were treated for 1 week with or without 1μM Fluoxetine hydrochloride (Sigma, #F132) in complete growth medium with 10% fetal bovine serum. Control and Fluoxetine conditions were tested in triplicate, with treatment/medium replacement every 48h. At the end of the treatment, total RNA was isolated with a Qiagen RNeasy kit. RNA concentration and integrity was assessed with a DS-11 Spectrophotometer (Denovix, Wilmington, DE). 50ng of RNA were retro-transcripted with a Maxima H Minus cDNA Synthesis Master Mix kit (Thermo). Expression of HTR2B and SLC6A4 and in response to Fluoxetine treatment was measured by TaqMan gene expression analysis on a Piko Real real-time PCR apparatus (Thermo) using FAM-based probes (HTR2B, Hs00168362_m1; SLC6A4, Hs00984349_m1, α-smooth muscle actin (Hs00426835_g1, Thermo) and vimentin (Hs00958111_m1, Thermo) were used as activation markers. GAPDH (Hs99999905_m1, Thermo) was used as housekeeping gene. Results were calculated with the 2-ΔΔCT method and expressed as a percentage of expression in untreated cells.

### Platelet Methodology

#### Blood Collection

Healthy human subjects on no medications were recruited with informed consent (Children’s Hospital of Philadelphia IRB Protocol #12-008608); see Tables S9 and S10 for demographics of the human subjects who participated in these studies Blood was collected using an 18 gauge needle into a tube preloaded with 4% w/v citrate buffer (cat# 68-04-2, SIGMA, St. Louis MO, USA).

#### Chandler Loop Studies

The Chandler Loop apparatus was employed, as previously described (1-3) to study platelet activation as a function of Fluoxetine or Aspirin. Aspirin, 10mM stock(cat # AS130, Spectrum, Gardena, CA, USA) was prepared, in 1M HEPES buffer pH 7.8. Fluoxetine (cat # 1279804, The United States Pharmacopeia, Rockville, MD, USA) was prepared, as a 1mM stock, in 1M HEPES buffer, at pH 7.8 and stored at −20C for experimental use. In brief, the procedure used was as follows: 10.0 ml of blood, acquired from donors described above, were treated with or without Fluoxetine (10 µM) or Aspirin (250 µM) and carefully injected into the Chandler Loop, composed of polyvinyl chloride tubing (TERUMO-CVS, Ashland MA, USA) with the following dimensions: 40 cm length, inner diameter of 6.35 mm, wall thickness of 1.59 mm). The Loop was rotated at 50 rpm, with a calculated shear force of 18 dynes/cm^2^, for 4 hours at 37C. At the conclusion of the study, the blood was removed from the tubing and processed as detailed below. For 5HT studies (below) blood samples were centrifuged at 200g at 24C for 20 minutes. Plasma was collected and stored at −20C.

#### Flow Cytometry

Blood was collected from the sample port of the Chandler Loop in 50 ml tubes to avoid spillage. 1ml samples of blood were pipetted into 15ml tubes pre-loaded with 0, 0.5, 2.0, and 5.0µM Adenosine diphosphate (ADP), a P2Y1, P2Y12, and P2X1 agonist (Catalog # 101312 Bio Data Corporation, Horsham, PA) and were incubated for 2 minutes at room temperature. 50 µl of ADP treated or non-treated blood were incubated with a monoclonal antibody specific for the platelet activation marker, P-Selectin, CD62P-labelled with R-phycoerythrin (PE) (P-selectin, clone AC1.2, cat# 348107, BD Bioscience, San Jose, CA, USA), and a monoclonal antibody for the platelet specific marker CD42b-labeled with allophycocyanin (APC) (GP 1BA, clone HIP1, cat# 551061, BD Bioscience). Post staining, platelets were fixed with 1% Formaldehyde (w/v) (cat# 28908, Thermo Fischer Scientific, Rockford, IL, USA) in the dark at room temperature for 15 minutes. Samples were analyzed using the BD AccuriTM C6 flow cytometer (BD Bioscience). A platelet gate was set based on the intensity of APC (CD42b) and activation was determined by the intensity of APC in conjunction with the intensity of PE (CD62P) and 300,000 events were recorded.

#### 5HT Measurements

5HT levels were determined by the Metabolomics Core Facility of The Children’s Hospital of Philadelphia Research Institute, using Agilent 1260 Infinity Triple Quad 6410B mass spectrometry (MS) coupled with LC. Isotope dilution methodology approach was used for 5HT measurement in plasma samples as follows:10 pmoles of D4-5HT (Adrich, Cat#673455, 5HT-α,α,β,β,-D4-creatinine sulfate monohydrate), were added to 150 uL of plasma sample, mixed with 140 uL of added PBS to bring the total volume of each sample to 300 µl Then, 160 µL of ethanol and 40 µL of pyridine was added followed by addition of 50ul ethyl chloroformate to derivatize 5HT in the sample., Samples were then gently shaken until bubbles disappeared. After shaking, 0.5 ml of water was added, and derivative was extracted twice with ethyl acetate by vigorous vortexing for 1 minute and span down. The upper organic phase was transferred to a clean vial and evaporated completely under air stream at 37oC. The dry sample was reconstituted in 100 µL of 0.1% formic acid in water. 5 µl of the derivatized sample was injected into the Agilent LC/MS/MS system. We used ions at m/z m/z 321-203 MRM for 5HT; and 325-207 MRM for D4-5HT. The concentrations of 5HT in the plasma were calculated by the isotope-dilution approach as previously described (4).

---

**Table S1.** Multivariate analysis: Cox Hazard Model Analysis of the 225 Patient Population with Surgery for Degenerative Mitral Valve Regurgitation

|  | Numbers of subjects N=225 |  | Age at Surgery (mean, years) |  | Hazard Ratio | Lower 95% confidence limit | Upper 95% confidence limit | p Value |
| --- | --- | --- | --- | --- | --- | --- | --- | --- |
| **Co-variate** | Present | % | no | yes |  |  |  |  |
| **Male** | 160 | 71.1% | 63.8 | 62.9 | 1.59 | 1.16 | 2.19 | 0.0035 |
| Aortic valve disease | 28 | 12.4% | 61.5 | 74.8 | 0.46 | 0.28 | 0.72 | <0.001 |
| Aspirin | 91 | 40.4% | 61.1 | 66.1 | 0.67 | 0.50 | 0.89 | 0.0055 |
| Atrial fibrillation | 52 | 23.1% | 62.0 | 67.0 | 0.68 | 0.49 | 0.94 | 0.0187 |
| Coronary artery disease | 79 | 35.1% | 59.3 | 70.3 | 0.49 | 0.35 | 0.68 | <0.001 |
| Heart Failure (NYHA) level | 93 | 41.3% | 60.9 | 66.3 | 0.68 | 0.50 | 0.90 | 0.0080 |
| Hyperlipidemia | 92 | 40.9% | 61.3 | 65.8 | 0.68 | 0.51 | 0.91 | 0.0088 |
| Renal insufficiency | 16 | 7.1% | 62.5 | 71.5 | 0.43 | 0.23 | 0.73 | 0.0013 |
| SSRI | 20 | 8.9% | 63.5 | 59.9 | 1.86 | 1.12 | 2.94 | 0.0183 |

**Table S2.** Multivariate Analysis Using the Optum National Database with a Cox Hazard Model Analysis of the Patient Population with Surgery for Mitral Valve Regurgitation Excluding those with Concomitant Coronary Artery Bypass Surgery

|  | Number of subjects<br>N=9,441 |  | Age at Surgery<br>(mean, years) |  | Hazard Ratio | Lower 95% confidence limit | Upper 95% confidence limit | p-value |
| --- | --- | --- | --- | --- | --- | --- | --- | --- |
| **Co-variate** | present | % | no | yes |  |  |  |  |
| Male | 5353 | 56.7% | 65.0 | 61.9 | 1.27 | 1.19 | 1.37 | <0.001 |
| Alcohol Dependence | 274 | 2.9% | 63.1 | 62.8 | 1.32 | 1.16 | 1.48 | <0.001 |
| Aortic Valve Disease | 2626 | 27.8% | 62.4 | 64.9 | 0.90 | 0.86 | 0.94 | <0.001 |
| Atrial Fibrillation | 3563 | 37.7% | 60.4 | 64.9 | 0.72 | 0.69 | 0.75 | <0.001 |
| Coronary Artery Disease | 4621 | 49.0% | 59.5 | 66.8 | 0.79 | 0.76 | 0.83 | <0.001 |
| Chronic Lung Disease | 2973 | 31.5% | 61.8 | 66.0 | 0.98 | 0.94 | 1.03 | 0.443 |
| Congestive Heart Failure | 4225 | 44.8% | 60.5 | 66.3 | 0.87 | 0.83 | 0.91 | <0.001 |
| Hypertension | 6433 | 68.1% | 57.2 | 65.8 | 0.70 | 0.66 | 0.74 | <0.001 |
| Obesity | 1146 | 12.1% | 64.3 | 62.1 | 1.42 | 1.33 | 1.51 | <0.001 |
| Peripheral Vascular Disease | 2292 | 24.3% | 61.3 | 68.7 | 0.75 | 0.71 | 0.78 | <0.001 |
| Psychiatric Disorder | 605 | 6.4% | 63.1 | 63.4 | 1.04 | 0.96 | 1.15 | 0.315 |
| Renal insufficiency | 1110 | 11.8% | 62.2 | 69.6 | 0.80 | 0.75 | 0.84 | <0.001 |
| SSRI | 604 | 6.4% | 64.1 | 62.3 | 1.28 | 1.19 | 1.37 | <0.001 |
| Stroke | 382 | 4.1% | 62.9 | 68.3 | 0.86 | 0.77 | 0.94 | 0.009 |

**Table S3:** 5HT Related Gene Expression Patterns Comparing Mitral Regurgitation (MR) Leaflets (44) versus Normal (20): Quantitative RT-PCR Results Showing Two Fold or Greater Differences

| Gene symbol | Description | Fold change | p |
| --- | --- | --- | --- |
| ADCY2 | Adenylate cyclase 2 | +2.38 | 0.003 |
| ADCY3 | Adenylate cyclase 3 | +3.48 | <0.001 |
| ADRB2 | Adrenergic, beta-2-, receptor | -2.15 | <0.001 |
| DDC | Dopamine decarboxylase | -5.99 | 0.005 |
| DRD3 | Dopamine receptor D3 | -5.29 | 0.001 |
| EPHB1 | EPH receptor B1 | -1.95 | 0.002 |
| HTR1B | 5HT receptor 1B | -5.52 | 0.002 |
| HTR1D | 5HT receptor 1D | -3.08 | 0.003 |
| HTR1E | 5HT receptor 1E | -4.19 | <0.001 |
| HTR1F | 5HT receptor 1F | -2.50 | 0.001 |
| HTR2C | 5HT receptor 2C | -3.89 | 0.031 |
| HTR3A | 5HT receptor 3A | -4.19 | <0.001 |
| HTR3B | 5HT receptor 3B | -4.58 | 0.003 |
| HTR5A | 5HT receptor 5A | -4.16 | <0.001 |
| HTR7 | 5HT receptor 7 | -3.25 | <0.001 |
| MAOB | Monoamine oxidase-B | +2.13 | 0.001 |
| NR4A3 | Nuclear receptor subfamily 4, group A, member 3 | -2.26 | 0.041 |
| PDE4B | Phosphodiesterase 4B | -3.13 | <0.001 |
| PDE4D | Phosphodiesterase 4D | -3.84 | <0.001 |
| PDYN | Prodynorphin | -4.37 | <0.001 |
| PIK3CA | Phosphoinositide-3-kinase, catalytic, alpha polypeptide | -13.73 | <0.001 |
| PIK3CG | Phosphoinositide-3-kinase, catalytic, gamma polypeptide | -4.31 | <0.001 |
| PLA2G5 | Phospholipase A2, group V | -3.19 | <0.001 |
| SLC18A1 | Vesicular monoamine transporter 1 | -3.94 | <0.001 |
| SLC18A2 | Vesicular monoamine transporter 2 | -18.57 | <0.001 |
| SLC6A3 | Dopamine transporter | -3.00 | 0.007 |
| SLC6A4 | 5HT transporter | -5.22 | <0.001 |
| SNCA | Synuclein, alpha | -21.21 | <0.001 |
| SYN2 | Synapsin II | -2.43 | 0.001 |
| TH | Tyrosine hydroxylase | -3.75 | 0.005 |
| TPH1 | Tryptophan hydroxylase 1 | -2.31 | 0.015 |

**Table S4:**
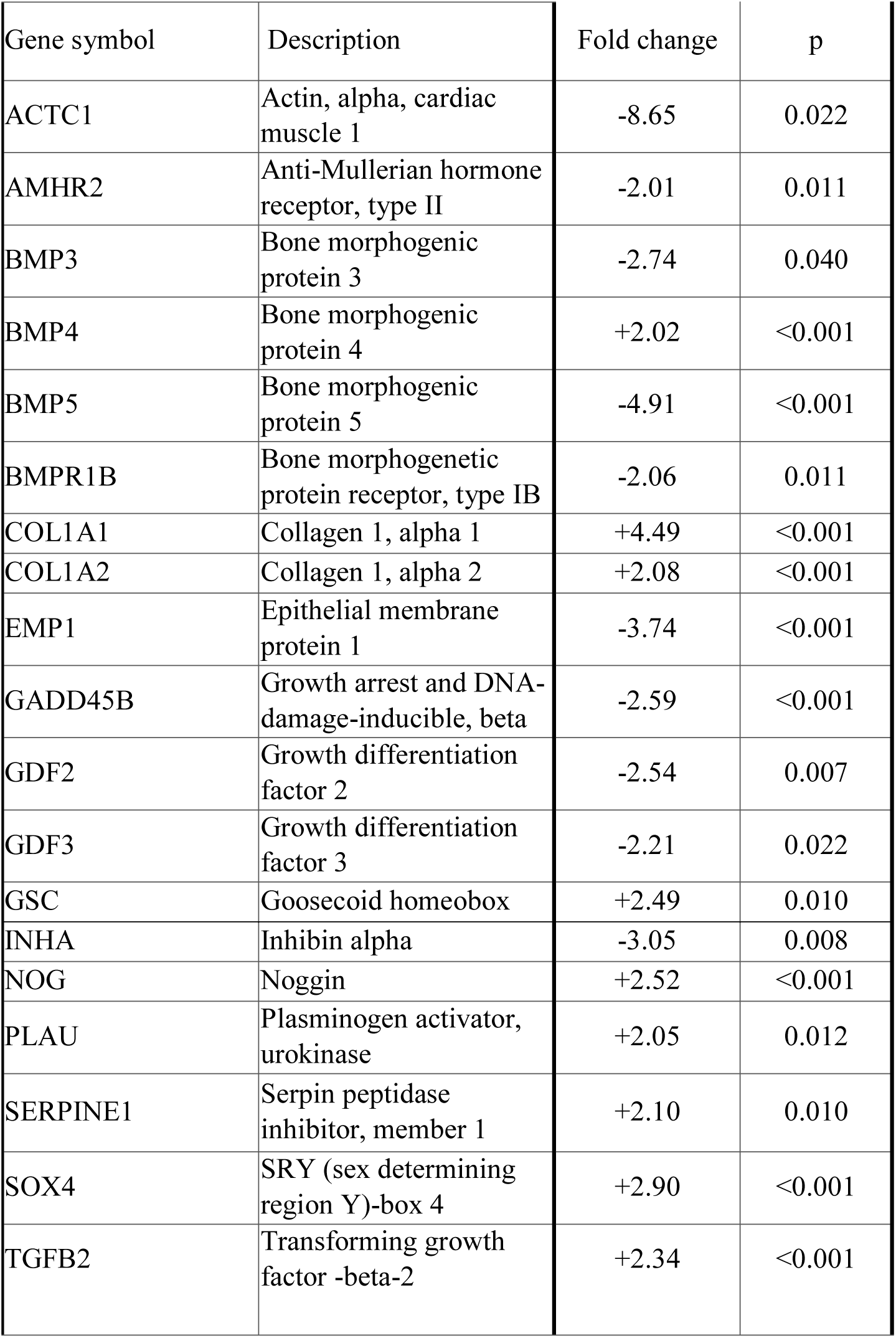
TGFβ Gene Expression Patterns Comparing Mitral Regurgitation (MR) Leaflets (40) versus Normal (19): Quantitative RT-PCR Results Showing Two Fold or Greater Differences

| Gene symbol | Description | Fold change | p |
| --- | --- | --- | --- |
| ACTC1 | Actin, alpha, cardiac muscle 1 | -8.65 | 0.022 |
| AMHR2 | Anti-Mullerian hormone receptor, type II | -2.01 | 0.011 |
| BMP3 | Bone morphogenic protein 3 | -2.74 | 0.040 |
| BMP4 | Bone morphogenic protein 4 | +2.02 | <0.001 |
| BMP5 | Bone morphogenic protein 5 | -4.91 | <0.001 |
| BMPR1B | Bone morphogenetic protein receptor, type IB | -2.06 | 0.011 |
| COL1A1 | Collagen 1, alpha 1 | +4.49 | <0.001 |
| COL1A2 | Collagen 1, alpha 2 | +2.08 | <0.001 |
| EMP1 | Epithelial membrane protein 1 | -3.74 | <0.001 |
| GADD45B | Growth arrest and DNA-damage-inducible, beta | -2.59 | <0.001 |
| GDF2 | Growth differentiation factor 2 | -2.54 | 0.007 |
| GDF3 | Growth differentiation factor 3 | -2.21 | 0.022 |
| GSC | Goosecoid homeobox | +2.49 | 0.010 |
| INHA | Inhibin alpha | -3.05 | 0.008 |
| NOG | Noggin | +2.52 | <0.001 |
| PLAU | Plasminogen activator, urokinase | +2.05 | 0.012 |
| SERPINE1 | Serpin peptidase inhibitor, member 1 | +2.10 | 0.010 |
| SOX4 | SRY (sex determining region Y)-box 4 | +2.90 | <0.001 |
| TGFB2 | Transforming growth factor -beta-2 | +2.34 | <0.001 |

**Table S5:** Characteristics of the mitral regurgitation subjects providing mitral valve interstitial cells

| Parameter | N |
| --- | --- |
| Total number | 5 |
| Men | 3 |
| Women | 2 |
| Age range (years) | 45-59 |

**Table S6:** Characteristics of the transplant subjects with normal mitral valves providing mitral valve interstitial cells

| Parameter | N |
| --- | --- |
| Total number | 5 |
| Men | 2 |
| Women | 3 |
| Age range (years) | 22-78 |

**Table S7:** Characteristics of the subjects participating in the Fluoxetine platelet study

| Parameter | N |
| --- | --- |
| Total number | 5 |
| Men | 2 |
| Women | 3 |
| Age range (years) | (24-33) |

**Table S8:**
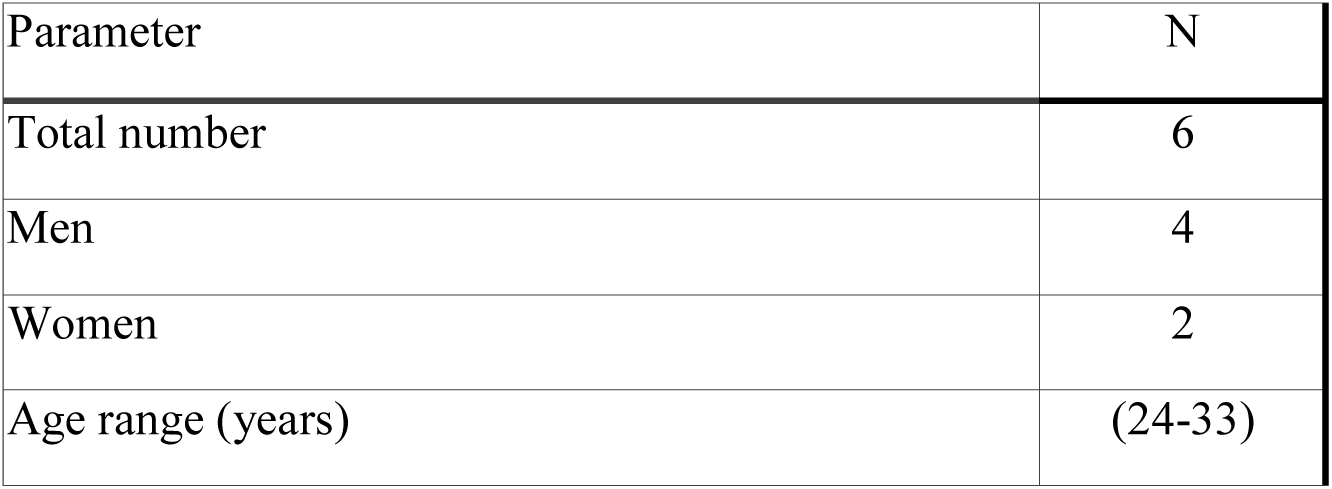
Characteristics of the subjects participating in the Aspirin platelet study

| Parameter | N |
| --- | --- |
| Total number | 6 |
| Men | 4 |
| Women | 2 |
| Age range (years) | (24-33) |

